# Get to know your neighbors with a SNAQ^TM^: A framework for single cell spatial neighborhood analysis in immunohistochemical images

**DOI:** 10.1101/2024.08.04.606539

**Authors:** Aryeh Silver, Avirup Chakraborty, Avinash Pittu, Diana Feier, Miruna Anica, Illeana West, Matthew R. Sarkisian, Loic P. Deleyrolle

## Abstract

**Motivation:** Analyzing the local microenvironment of tumor cells can provide significant insights into their complex interactions with their cellular surroundings, including immune cells. By quantifying the prevalence and distances of certain immune cells in the vicinity of tumor cells through a neighborhood analysis, patterns may emerge that indicate specific associations between cell populations. Such analyses can reveal important aspects of tumor-immune dynamics, which may inform therapeutic strategies. This method enables an in-depth exploration of spatial interactions among different cell types, which is crucial for research in oncology, immunology, and developmental biology.

**Results:** We introduce an R Markdown script called SNAQ^TM^ (**<u>S</u>**ingle-cell Spatial **<u>N</u>**eighborhood **<u>A</u>**nalysis and **<u>Q</u>**uantification), which conducts a neighborhood analysis on immunofluorescent images without the need for extensive coding knowledge. As a demonstration, SNAQ^TM^ was used to analyze images of pancreatic ductal adenocarcinoma. Samples stained for DAPI, PanCK, CD68, and PD-L1 were segmented and classified using QuPath. The resulting CSV files were exported into RStudio for further analysis and visualization using SNAQ^TM^. Visualizations include plots revealing the cellular composition of neighborhoods around multiple cell types within a customizable radius. Additionally, the analysis includes measuring the distances between cells of certain types relative to others across multiple regions of interest.

**Availability and implementation:** The R Markdown files that comprise the SNAQ^TM^ algorithm and the input data from this paper are freely available on the web at https://github.com/AryehSilver1/SNAQ.

**Visual Abstract:** 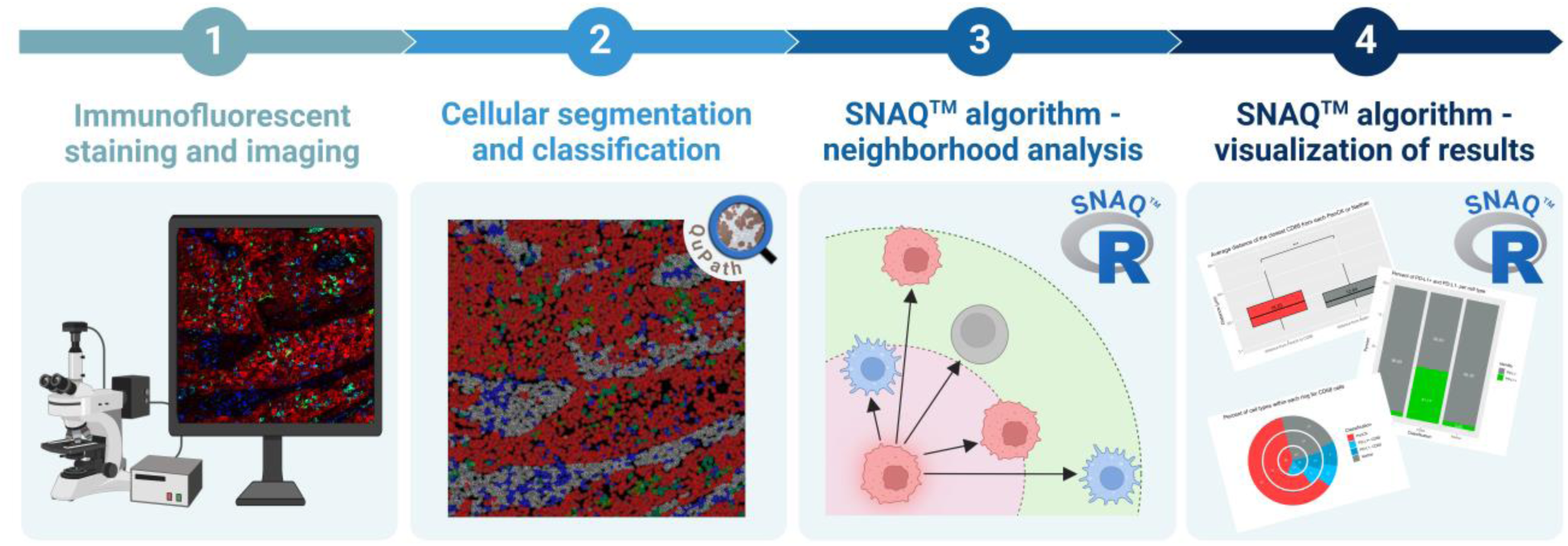

Created with BioRender.com.

## 1. Introduction

Exploring the highly complex and heterogeneous ecosystem of the tumor microenvironment (TME) provides valuable insights into the intricate interactions among tumor cells, stromal tissues/cells, the extracellular matrix, and the immune microenvironment. This aids in forming a comprehensive overview and detailed insight at the spatial-molecular resolution into the dynamics between the cells, providing potential insights into the cellular interactions that influence tumor behavior (Jia *et al*. 2022). Our proposed method for neighborhood analysis facilitates the quantitative assessment of spatial relationships within the complex tumor microenvironment. This involves analyzing how specific cell types are distributed and aligned in relation to tumor cells, assessing both their prevalence and proximity; thus, providing a more detailed understanding of the overall picture of the cellular interactions within the TME from a single snapshot. This approach provides details into the spatial dynamics crucial for understanding interactions within tumors and their surrounding immune milieu, offering important insights into spatial organization and uncovering distinct spatial patterns in tumor samples. These insights are pivotal to better comprehend the critical aspects of tumor-immune cell (any cell subtype) dynamics, which may form the basis of targeted therapeutic interventions to manipulate immune responses leading to efficacious cancer therapy (Jia *et al*. 2022).

While open-access image analysis tools like QuPath (https://qupath.github.io/) and CellProfiler (https://cellprofiler.org/) facilitate high-throughput image analysis leading to identification of cell types and quantification of biomarker expression of immunohistochemically stained tissue samples, they have limited capabilities for conducting neighborhood analyses based on cell type. While QuPath can measure properties of classified objects, such as the distance to the nearest object or annotation of a given class for each cell, it does not provide detailed information about the number and types of neighboring cells within specific distances from cells of a particular class. Additionally, CellProfiler does not support the execution of comprehensive geospatial analyses.

This study introduces an analysis pipeline for creating and executing custom neighborhood analyses on immunofluorescent tissue samples using RStudio post QuPath cell segmentation and classification. The algorithm, named SNAQ^TM^ (**<u>S</u>**ingle-cell Spatial **<u>N</u>**eighborhood **<u>A</u>**nalysis and **<u>Q</u>**uantification), facilitates the identification and quantification of cell types in proximity to any specified cell type. It also allows for the visualization and measurement of distances between different cell types. By employing this algorithm, valuable insights into the spatial relationships and interactions within the tissue microenvironment can be obtained, enhancing the understanding of cellular organization and behavior in various biological contexts. The code is highly customizable, allowing users to modify specifics to fit their unique image sets. The algorithm has an automated workflow and allows for batch processing of multiple images, empowering users with minimal coding experience to perform neighborhood analyses on their tissues. While this study demonstrates an example of using our neighborhood algorithm, it is designed for the research community to apply to various tissues and marker selections. Thus, to demonstrate its universal applicability, instead of hardcoding marker names in the algorithm, we anonymized them by assigning letter codes that are used consistently across the R Markdown scripts, as detailed in **Table 1**.

**Table 1.** Summary of each marker and corresponding cell type. The “Letter Code” column lists the letter codes that replace the marker names in the RStudio scripts to facilitate customization. The “Marker Meaning” column provides the ideal representation that each letter code should denote, guiding marker selection. The “Markers Used” and “Classification” columns are specific to this paper and can be modified to accommodate different markers and resulting classifications. Notably, since DAPI is not included in the code, this channel has not been assigned a letter.

| Letter Code | Marker Meaning | Markers Used | Classification |
| --- | --- | --- | --- |
| -- | Nuclear stain | DAPI | Nuclei |
| A | Cell Type 1 | PanCK | Tumor cells |
| B | Cell Type 2 | CD68 | Macrophages |
| C | Functional marker for B | PD-L1 | Immunosuppression marker |
| D | Non-Cell Type 1 + Non-Cell Type 2 | Negative for both PanCK and CD68 | Neither tumor cell nor macrophage |

The neighborhood analysis consists of two components: quantifying the number of neighbors within specified distances from each cell and finding the closest neighbor of a certain classification for each cell type. The former requires the visualization of concentric rings around each cell, where the number of cells that lie within each ring, as well as their classification, are recorded. The concentric rings and their distances from the target cell are displayed in **Figure 1**. The comparison of the different concentric rings offers a tool to assess specific spatial relationships, evaluate cell proximity, and identify interaction zones to provide insights into the cellular architecture within the TME and potentially understand immune evasion to inform therapeutic strategies. The latter involves calculating the distance between the target cell and the closest cell of a specific type, such as a tumor cell or macrophage, providing quantitative metrics across different cell types within samples. The results of the neighborhood analysis can be visualized in multiple ways using R Markdown, revealing patterns that may provide valuable insights into the interactions between different cell types. Of note, in the current study, the SNAQ^TM^ algorithm placed greater emphasis on B cells that are positive for the C functional marker, as opposed to those that are negative for it. However, data on B cells negative for the C marker are still collected and can be analyzed if relevant.

**Figure 1.**
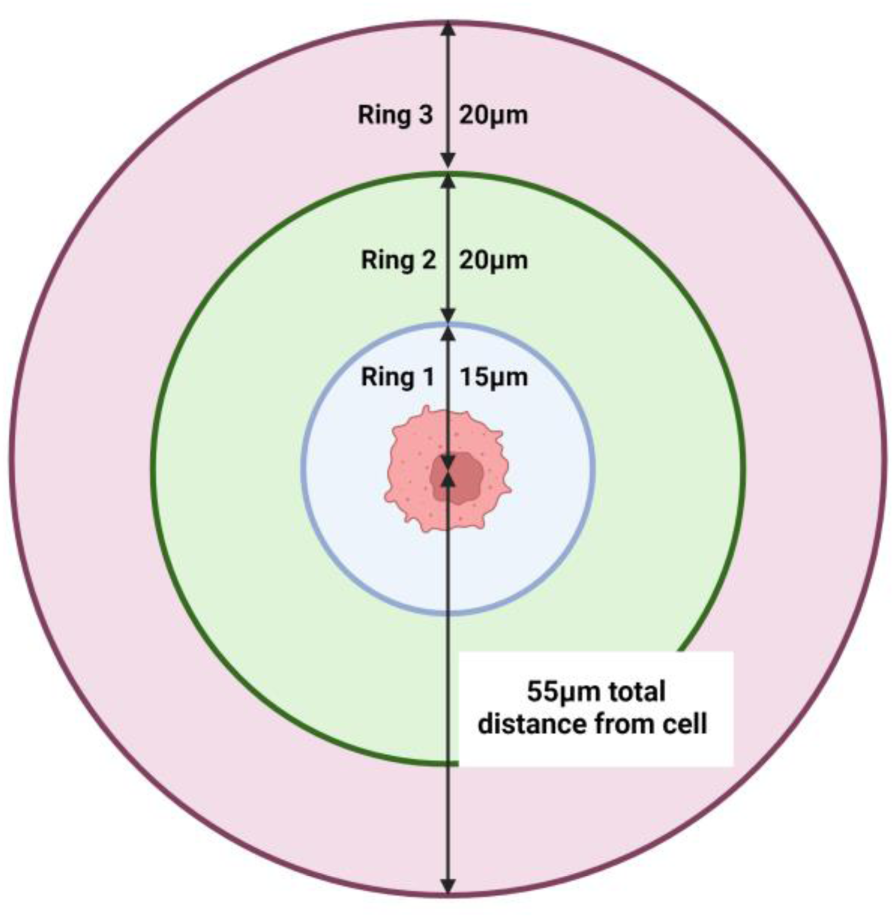
Diagram showing the concentric rings visualized around a target cell. The proximal neighborhood (Ring 1) forms the first compartment, closest to the central cell, extending 15 µm from the center of the target cell (706.86 µm^2^). The intermediate neighborhood (Ring 2, 3141.59 µm^2^) extends 20 µm beyond the edge of Ring 1, and the distal neighborhood (Ring 3, 5654.87 µm^2^) extends an additional 20 µm from the edge of Ring 2. Thus, the total distance from the center of the target cell to the outer edge of the distal ring is 55 µm, defining the entire neighborhood. The area of the entire neighborhood is 9503.32 µm^2^.Created with BioRender.com.

Pancreatic ductal adenocarcinoma (PDAC) is an immunologically cold tumor that is characterized by substantial infiltration of tumor-associated macrophages (TAMs), which are the most common infiltrating immune cell in the tumor microenvironment (Karamitopoulou 2019; Kane, Pa-C *et al*. 2020). These activated TAMs contribute to desmoplasia and are a poor prognostic indicator (Poh and Ernst 2021). Through stimulation of the PD-1/PD-L1 axis, macrophages can be polarized towards the M2 phenotype (Wei *et al*. 2021). M2-like macrophages contribute to the immunosuppressive TME characteristic of cold tumors by expressing PD-L1, which can inhibit the activation of cytotoxic T cells and helps create an immune-privileged environment (Pratt *et al*. 2021). Better characterization of the geospatial relationship between tumor-associated macrophages and tumor cells in PDAC may help guide novel therapies.

To demonstrate the capabilities of our algorithm, we used immunofluorescent scans of human pancreatic ductal adenocarcinoma (PDAC) (Aleynick *et al*. 2023). The scans contains the following markers and fluorophores: 4’,6-diamidino-2-phenylindole (DAPI) as a nuclear stain, Cy7 for pan-cytokeratin (PanCK), Cy3 for CD68, and Cy5 for programmed death-ligand 1 (PD-L1). PanCK is used as a tumor cell marker (Menz *et al*. 2023), and CD68, a widely recognized myeloid cell marker, identifies macrophages (Chistiakov *et al*. 2017). PD-L1, which binds to PD-1 on T cells, inhibits their proliferation, survival, and effector functions, suppressing the immune response against tumors (Han, Liu and Li 2020). TAMs express PD-L1, correlating with decreased survival in adenocarcinoma (Shinchi *et al*. 2022). In this study, we define macrophages expressing PD-L1 as immunosuppressive. **Table 1** outlines the markers and their corresponding cell types, while **Figure 2** illustrates the classification logic. For our analysis, PanCK and CD68 were used for cell typing, while PD-L1 served as a functional marker for macrophages. A lettering system has been implemented to represent these markers in the code, facilitating customization. The nuclear stain DAPI is necessary for cell segmentation in QuPath. Two markers are required for cell typing, each identifying a unique cell type. PanCK and CD68 classified cells as tumor cells, macrophages, or neither (**Figure 2**). Additionally, a functional marker is needed to sub-describe one of the cell types; PD-L1 was used to determine macrophages’ immunosuppressive status (**Figure 2)**. The functional marker must modify the marker represented by letter code B, which means that markers represented by letter codes A and B cannot be used interchangeably.

**Figure 2.**
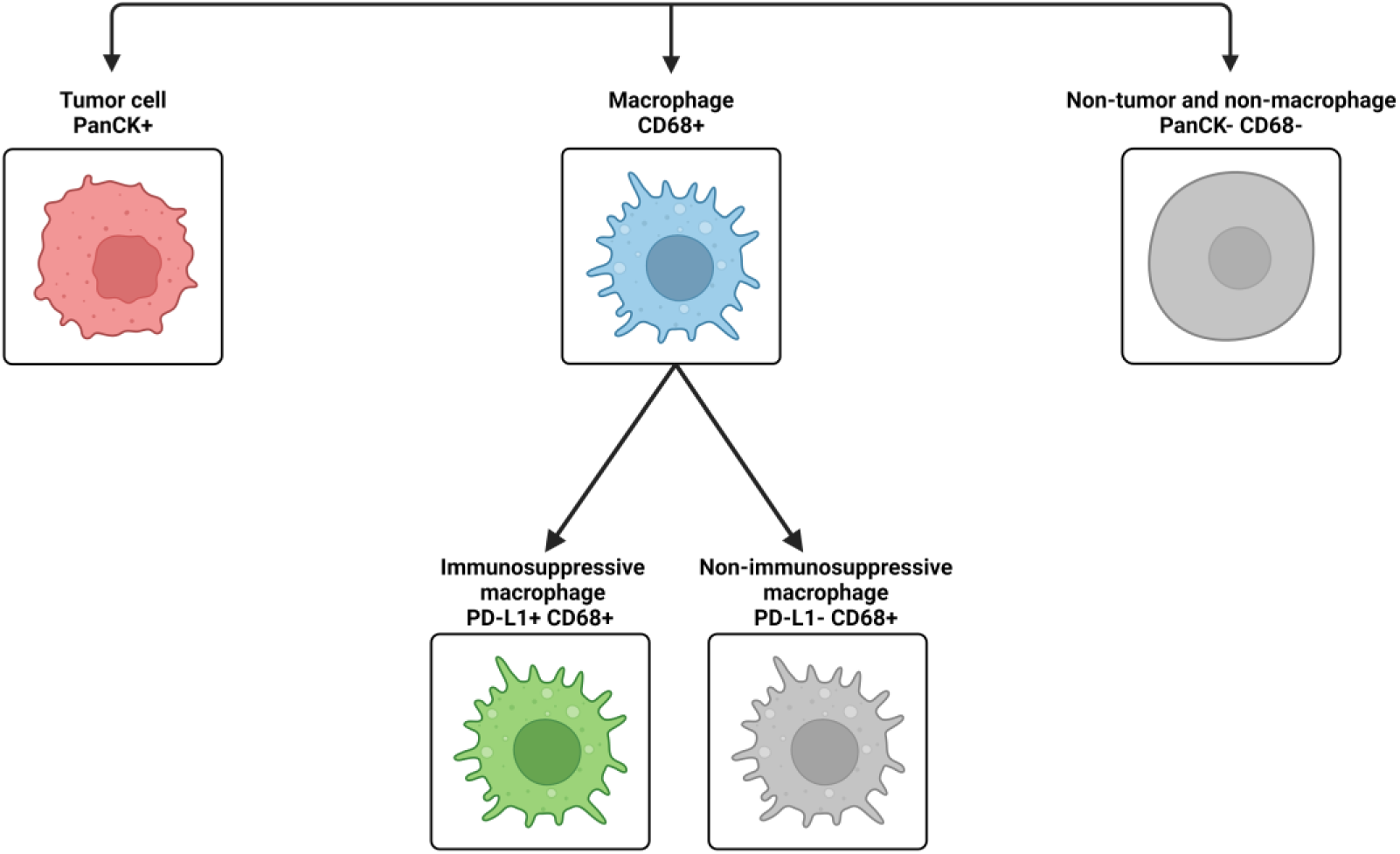
Classification schematic used in the PDAC image to label each cell. Both the phenotypic description and the markers are included. The top row is a cell’s classification based on expression of PanCK and CD68 and is used for cell typing, and the bottom row is a macrophage’s functional classification based on expression of PD-L1. Created with BioRender.com.

## 2. Methods

### 2.1. Image Acquisition

A large tile stitch of a PDAC tissue sample titled `PDAC(35000,27720)6800,3050` was accessed from a publicly-available dataset released under a Creative Common CC BY 4.0 license by Aleynick and colleagues (Aleynick *et al*. 2023). The stitch was acquired with a Zeiss Axioscan at a resolution of 0.3250 µm/pixel (Aleynick *et al*. 2023). The four markers utilized in this cellular neighborhood analysis study are nuclei, PanCK, CD68, and PD-L1, captured with the following dye or fluorophores DAPI, Cy7, Cy3, and Cy5, respectively. Representative images from selected regions of interest are shown in **Figure 3** for each marker.

**Figure 3.**
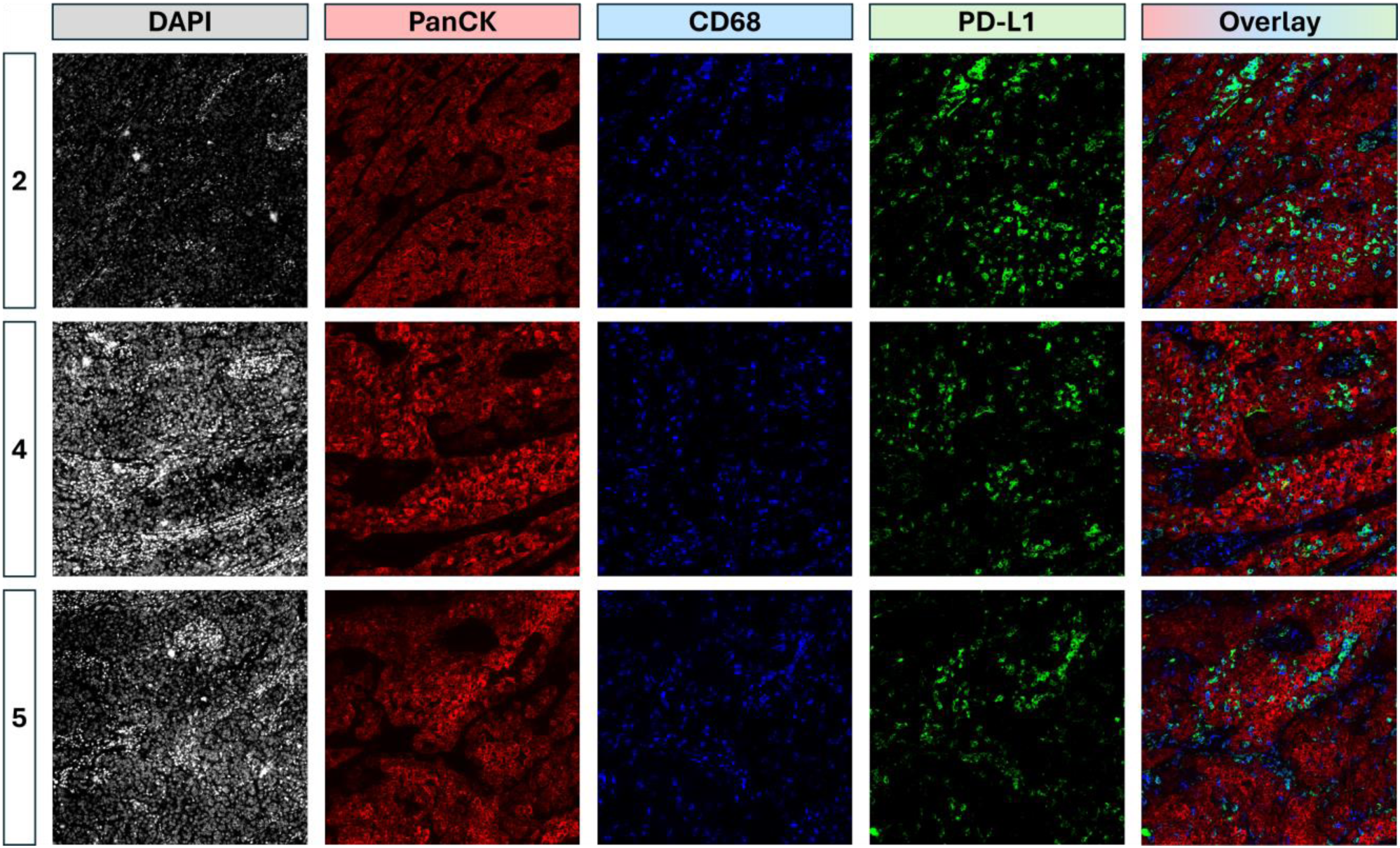
Representative images showing DAPI, PanCK, CD68, and PD-L1 markers. The images have been pseudocolored for visual contrast. The numbers on the left correspond to the ROI’s number, and shown are ROI_2, ROI_4, and ROI_5. Each ROI measures 750 µm by 750 µm. The Overlay image contains the markers PanCK, CD68, and PD-L1.

### 2.2. Cell Detection and Classification

Cellular detection and segmentation were achieved in QuPath v0.5.1 (Bankhead *et al*. 2017). A new project was created, and an annotation was drawn around the tumor area using the wand tool. Cell detection was run on the annotation based on the DAPI channel, and the parameters are shown in **Figure 4**. Five classes were created: PanCK, CD68, Neither, PD-L1, and Ignore*.

**Figure 4.**
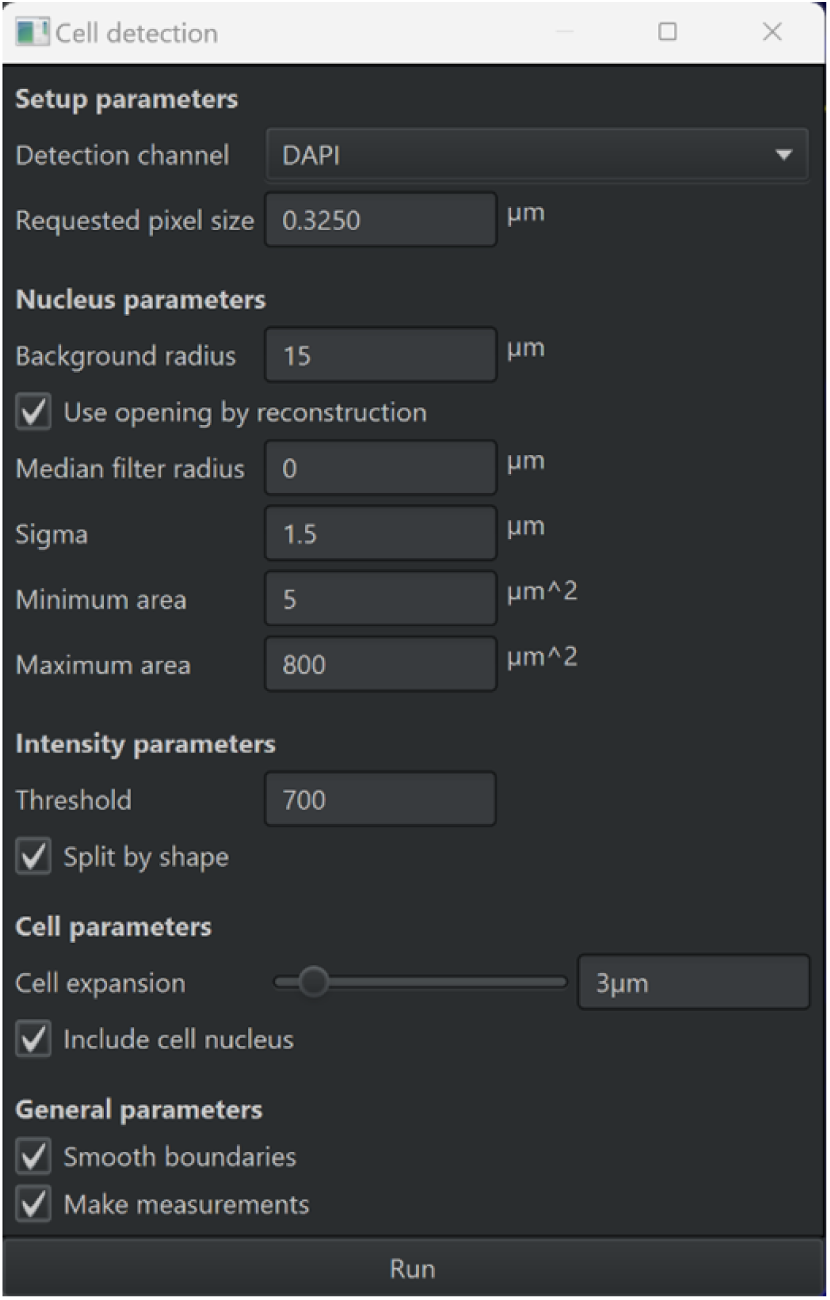
Parameters used for cell detection. Please note that the values of these parameters can be adjusted to optimize detection based on the type of tissue and the acquired image.

An object classifier that can distinguish between cells that are either tumor cells (PanCK+), macrophages (CD68+), or neither (PanCK- and CD68-) was created. The point annotation tool was used to label 45 PanCK+ cells, which were then assigned to the PanCK class. Subsequently, 45 CD68+ cells were labeled and assigned to the CD68 class. Finally, another 45 cells that were both PanCK- and CD68-were labeled and assigned to the Neither class. The `Train Object Classifier` tool was used to train an object classifier based on the point annotations. All default settings were maintained except for the `Features` setting, which was adjusted to only consider measurements related to PanCK and CD68. To evaluate the accuracy of the classifier, the `Load Training` option was selected. It is important to note that while this paper annotated 45 cells per class to train the object classifier, the number of annotations required may vary for different images to ensure an accurate classifier. Depending on the complexity and variability of the tissue samples, more or fewer annotations might be necessary to achieve reliable classification results. Adjusting the number of annotations based on the specific characteristics of the images being analyzed can significantly enhance the performance of the object classifier. The classifier was saved as `Classification`. Next, a single measurement classifier was created to determine which cells were PD-L1+ or PD-L1-. The parameters to create this classifier are shown in **Figure 5**. Cells that are PD-L1+ are assigned to the PD-L1 class, and cells that are PD-L1-are assigned to the Ignore* class. The nomenclature “Ignore” serves as a placeholder for the absence of classification for PD-L1 negative macrophages (i.e., non-immunosuppressive) specific to the current study. However, this classifier’s name can be edited if desired. Save the classifier as `FunctionalClassification`.

**Figure 5.**
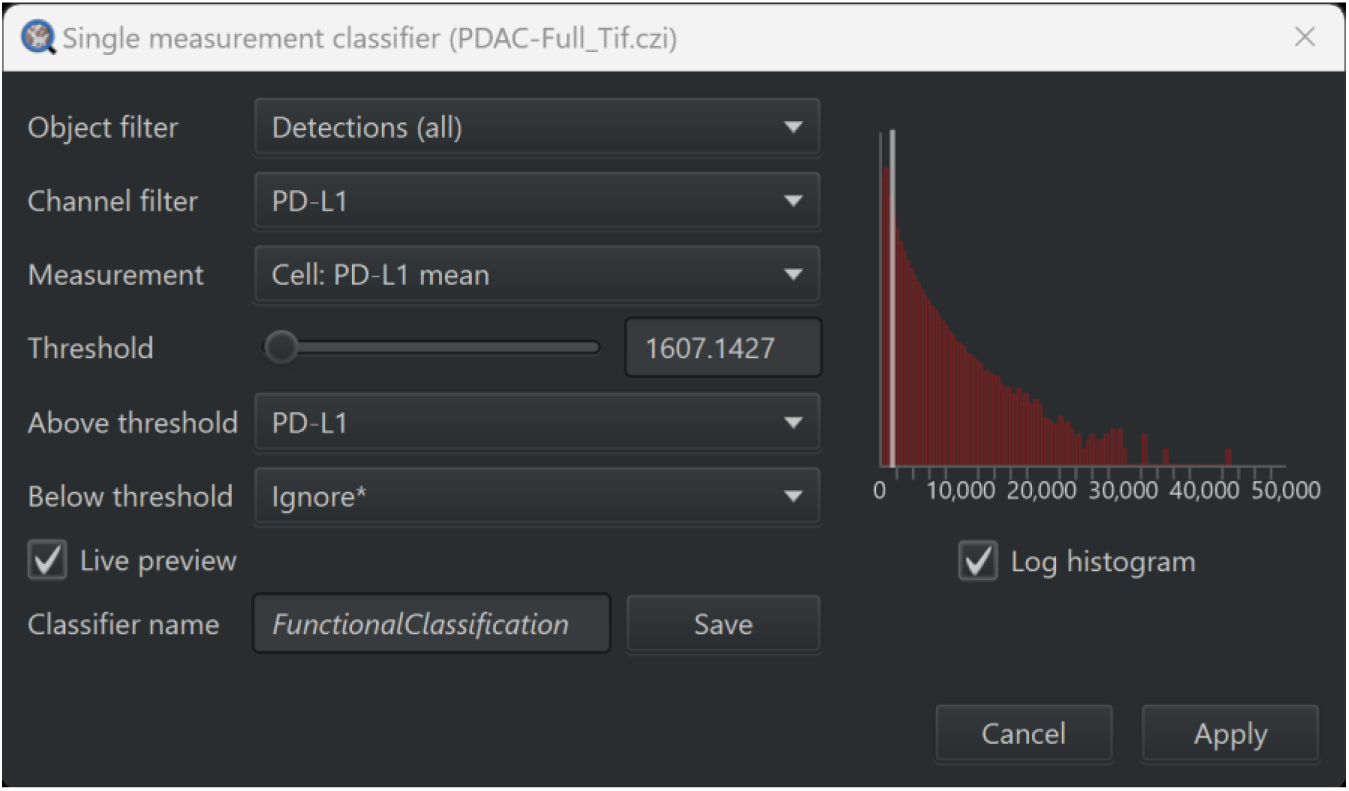
Parameters used to create the single measurement classifier for PD-L1. Note that the Threshold value needs to be optimized if a different image is used.

A composite classifier was created to combine `Classification` and `FunctionalClassification`, which was named `Combined` and applied to the image for the classification to each cell. Eight square annotations measuring 750 µm by 750 µm were scattered in random locations within the tumor annotation. Hierarchies were resolved with the shortcut Ctrl+Shift+R to insert the ROIs into their proper place in the hierarchy of annotations. The data are then exported as a CSV file, with the parameters for the cell data export shown in **Figure 6** and including Classification, Parent, Centroid X µm, and Centroid Y µm. The file saved as `PDAC_measurements.csv` should be placed in a folder specifically designated to hold the input data for the algorithm.

**Figure 6.**
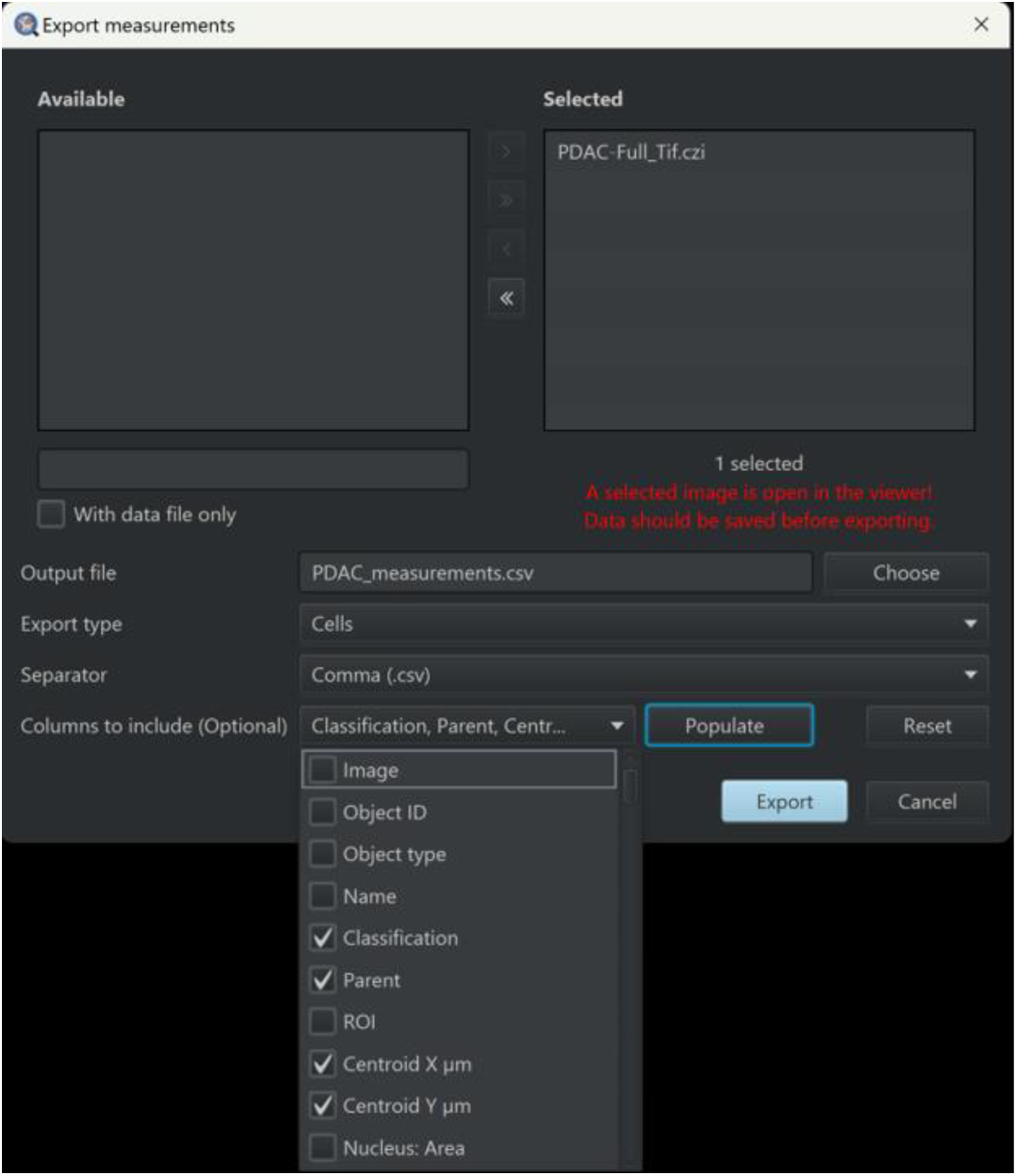
Parameters to export the data for the cells as a CSV file.

Representative images of the cell classification are shown in **Figure 7**, which shows both the Classification and FunctionalClassification classifiers.

**Figure 7.**
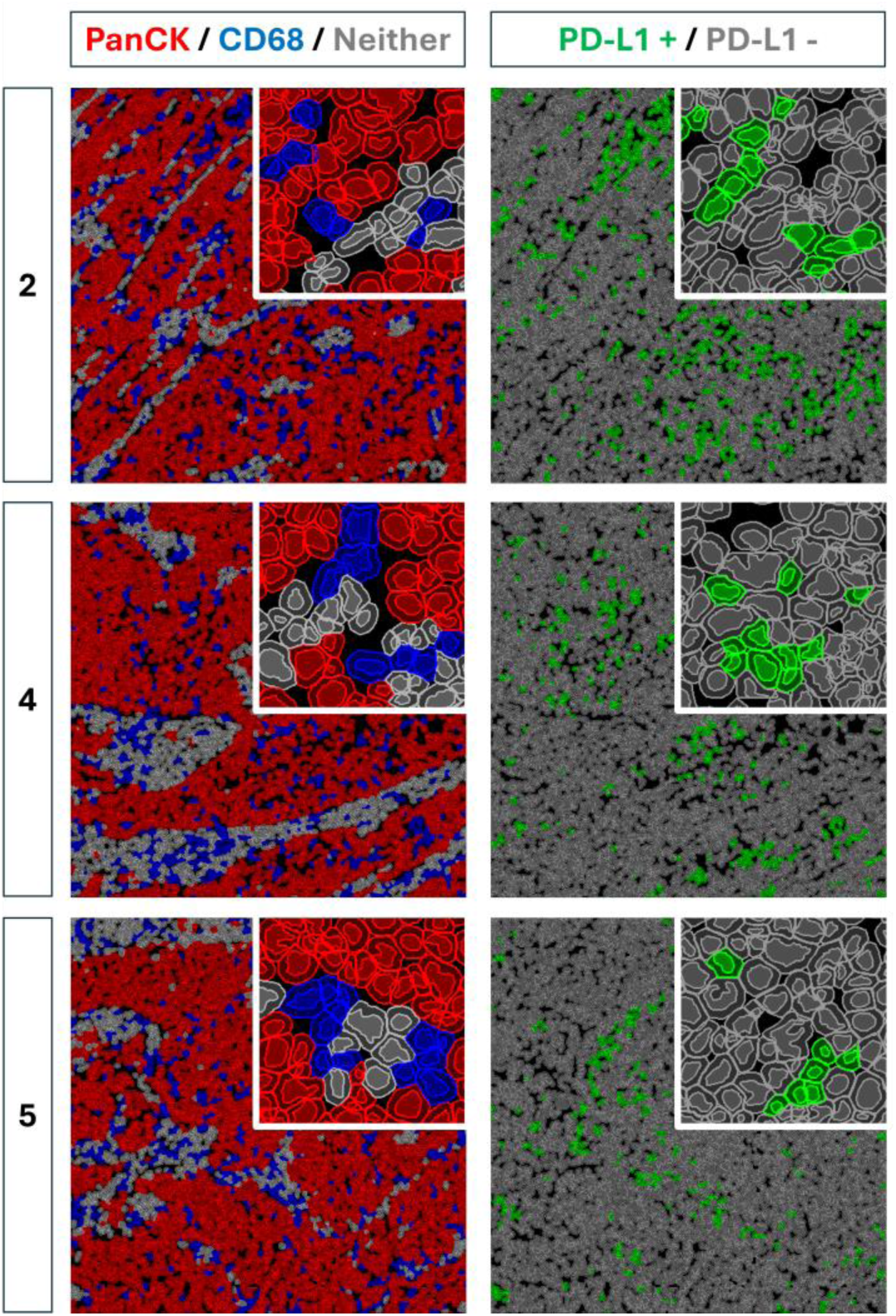
Representative images of the cell classification for ROI_2, ROI_4, and ROI_5. In the left panels, which represent the `Classification` classifier, cells are color-coded as follows: red for `PanCK` class, blue for `CD68` class, and grey for `Neither` class. For the column on the right, which represents the `FunctionalClassification` classifier, cells belonging to the PD-L1 class are green and cells belonging to the Ignore* class are grey.

### 2.3. Data Analysis

R Studio was employed to run the neighborhood analysis algorithm on the data derived from QuPath (Posit team 2024). The R Markdown file `Data Analysis.Rmd` was used to run the neighborhood analysis and exports the results for downstream visualization (Wickham *et al*. 2019; Microsoft and Weston 2022; Microsoft Corporation and Weston 2022; Dowle and Srinivasan 2023; Wickham and Bryan 2023; Arnold 2024). **Table 2** lists all the variables that need to be initialized within both the `Data Analysis.Rmd` and `Plot Maker.Rmd` R Markdown files. However, each file contains only a subset of these variables. Therefore, only the variables named in the “Variables to Initialize” code chunk in each respective file need to be assigned a value.

**Table 2.** List of all the variables that must be initialized before running the neighborhood analysis and plot maker. Also included is a description of what each variable represents and its data type. These variables are used across the **`Data Analysis.Rmd`** and **`Plot Maker.Rmd`** R Markdown files, and as such, each file will not contain all these variables. These variables must be set by the user to accommodate their images.

| Variable Name | Data Type | Description |
| --- | --- | --- |
| dist1 | Numeric | Desired thickness of Ring 1 in $\mu\text{m}$ |
| dist2 | Numeric | Desired thickness of Ring 2 in $\mu\text{m}$ |
| dist3 | Numeric | Desired thickness of Ring 3 in $\mu\text{m}$ |
| maxXum | Numeric | Length of each ROI in $\mu\text{m}$ |
| maxYum | Numeric | Height of each ROI in $\mu\text{m}$ |
| numImages | Numeric | Number of ROIs being analyzed |
| markerA | Character | Name of Marker A |
| markerB | Character | Name of Marker B |
| markerC | Character | Name of Marker C |
| inputFolder | Character | File path of the folder which contains the input CSV files |
| outputFolder | Character | File path of the folder to where the results will be saved |
| workingDir | Character | Same file path as `outputFolder` |
| graphOutputPath | Character | File path of the folder to where the data visualizations will be saved |

The master CSV file `PDAC_measurements.csv` was then imported into the `Data Analysis.Rmd` file. If the images were taken from a larger stitched image, such as when the ROIs were saved from a larger tile stitch of PDAC, then individual CSV files for each image must be created by filtering the master file. The centroid x- and y-axis positions for each ROI in the tile stitch were recorded, enabling the recalculation of the x- and y-coordinates for each cell so that the top left corner of each ROI becomes the origin. A separate CSV file was created for each ROI, containing only the cells that fall within it and their recalculated coordinates. These ROI-specific files are saved in the designated input folder.

The CSV files for each image were then aggregated into a master data frame called “`combo`.” During this process, cell numbers were reset for each new image, and an image number was assigned to each cell to keep track of its origin. The SNAQ^TM^ algorithm is designed to analyze one image at a time, which is achieved by wrapping the entirety of the main algorithm in a `for` loop that iterates through each image individually. For each iteration, `combo` is filtered to only include cells from the image currently being analyzed, and these cells are saved in a new data frame called `newCombo` that is rewritten after each iteration. For the remainder of this section, the process described occurred to each image separately.

In the data frame “`newCombo`,” the columns “`xDim`” and “`yDim`” store each cell’s x- and y-coordinates, respectively. A distance matrix is then calculated for each cell based on these coordinates. Each element in the matrix represents the distance between two cells, with the element’s row and column numbers corresponding to the “`ObjectNumber`” of the cells. Cells that are within 55 µm from the image border are excluded from the distance matrix calculation and return null values; however, cells that are not excluded still calculate a distance between themselves and excluded cells. The distance matrix is created using a `foreach` loop coupled with `%dopar%` to allow for parallel processing by distributing tasks to multiple workers, and this `foreach` loop is set equal to a master list called `distances` which combines the results from each worker. The use of multiple workers, representing independent units that concurrently perform specific computing tasks, allows for the simultaneous calculation of distances between cells and the appending of these values to the list `distances`. The `foreach` loop iterates through each cell in the image, and the cell focused on is called the target cell. A nested `for` loop finds the distance between the target cell and every other cell in the image (which is denoted as the secondary cell) through a custom function called `calcDist` that calculates Euclidean distance using `xDim` and `yDim` from both cells, with the formula for distance shown in **Formula 1**. `calcDist` takes as input two cells from `newCombo` and returns a numeric distance value in µm. The distance values for each target cell are returned as a list, which is concatenated to the master list `distances`. If the target cell is to be excluded, then the list that is returned is populated with null values and has a length equal to the number of cells in the image. Outside of the `for` loop, the master list `distances` is converted into a matrix to create the distance matrix (which is also named `distances`) that can be used to find the distance between any two cells.

**Formula 1.**
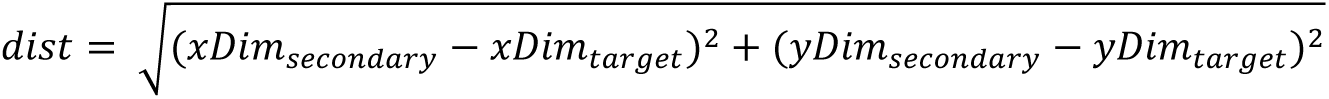
Euclidian formula used by the function calcDist to calculate the distance between two cells.

Note that when accessing the distance matrix `distances`, it is important to maintain the correct order of the indices. Since only non-excluded cells passed through the distance function, the x- and y-axes of the distance matrix are not interchangeable when retrieving values. The way to find the distance between a target cell and any secondary cell is through `distances[ObjectNumber` (of target cell)`, ObjectNumber` (of secondary cell)]. Reversing this order may return a null value if the location of the secondary cell falls within 55 µm from the image border and is subsequently excluded. This is especially important when iterating through the distances between a target cell and other secondary cells in the image. Another way to understand it is that the distance from the target cell to the secondary cell is determined by prioritizing the index of the target cell first. The distance matrix is used in conjunction with `newCombo` to calculate a neighborhood analysis for each cell. To spatially resolve the cell types at varying distances from the target cell, the number of neighbors within three concentric rings around the target cell are recorded. Ring1, defining the proximal neighborhood, spans from 0 µm to 15 µm away from the cell and provide information of the cell population in the immediate vicinity of the target cell. Ring 2, representing the intermediate neighborhood, covers distances from 15 µm to 35 µm. Finally, Ring 3, corresponding to the distal neighborhood, extends from 35 µm to 55 µm away from the cell. The counts from the three concentric rings (Ring 1, Ring 2, and Ring 3) can be aggregated to obtain the total number of neighboring cells within the entire neighborhood, spanning from 0 µm to 55 µm away from the target cell. The number of neighboring cells that are classified as tumor cells, macrophages, or neither are recorded, along with whether each neighbor is PD-L1 positive or negative. This data is then converted into proportions, reflecting the types of neighbors within the concentric rings as well as the total neighbors within the 55 µm radius environment. This was achieved using dynamic programming, where a `for` loop iterates through each cell in the image (which is the target cell), and a nested `for` loop iterates through every other cell (which is the secondary cell). Within the nested `for` loop, an `if` statement ensures that the target and secondary cells are not the same cell and that the value of the distance between the target and secondary cell is not a `null` value. Another `if` statement checks whether the distance between the target and secondary cells falls within a ring, and if it does, adds a value of one to the counter for the specific ring that corresponds to the secondary cell identity, PD-L1 status, and if applicable, its PD-L1+ macrophage cell status. These counts are used to tally the number and type of neighbors within each ring and keep a tally for the total number of neighbors across the three rings. Outside of the `for` loop and nested `for` loop, these counts are converted to proportions based on the totals for each ring, and the results are saved in columns in `newCombo`. `newCombo` is exported as a CSV file, with the image number being concatenated to the end of `newCombo`. For example, analysis of Image #2 exports newCombo2.csv. The distance matrix `distance` and the data stored in `newCombo` were used in combination to run a series of closest neighbor distance calculations, which are listed in **Table 3**.

**Table 3.** List of the closest neighbor calculations performed by the SNAQ^TM^ algorithm. The distances are calculated for each cell in the image that was not excluded due to proximity to the image border, and mean is used to find the average distance for each row. Only the data for the rows shaded in blue are displayed in the Results section as they have biological significance to the PDAC tissue.

| The distance of... |
| --- |
| the closest tumor cell from each macrophage |
| the closest neither cell from each macrophage |
| the closest tumor cell from each PD-L1+ macrophage |
| the closest neither cell from each PD-L1+ macrophage |
| the closest cell of any type from each macrophage |
| the closest cell of any type from each tumor cell |
| the closest cell of any type from each neither cell |
| the closest macrophage from each tumor cell |
| the closest macrophage from each neither cell |
| PD-L1+ macrophage from each tumor cell |
| the closest PD-L1+ macrophage from each neither cell |
| the 10 closest macrophages from each tumor cell |
| the 10 closest macrophages from each neither cell |

This was achieved by using a `foreach` loop with `%dopar%` that returns the shortest distance values for each target cell in the image and stores these values in a list. The target cell type is the one from which the measurements are being taken, and the secondary cell type is the cell type to which the target cell type is measured (e.g., distance of the closest macrophage from tumor cell, tumor cell is target cell type and macrophage is secondary cell type). `newCombo` is filtered to only include cells that are of the target cell type. A nested `for` loop iterates through every cell in the image, and an `if` statement ensures that only cells of the secondary cell type are taken for consideration. The distance between the target cell and the closest secondary cell of the specified cell type is found and concatenated to the list of shortest distance values. This list is exported as a CSV file to allow for downstream statistical analysis, and a separate list is exported for each type of measurement being recorded.

### 2.4. Data Visualization

The output from the neighborhood analysis was imported into the R Markdown file `Plot Maker.Rmd` (Grantham 2019; Wickham *et al*. 2019; Ahlmann-Eltze and Patil 2021; Dowle and Srinivasan 2023; Wickham and Bryan 2023). The variables that were initialized are listed in **Table 2**. The data from each image were compiled into master data frames for data visualization and statistical analysis.

## 3. Results of SNAQ^TM^

### 3.1. Cell Type Distribution and PD-L1 Expression in PDAC Microenvironment

Our analysis provides a breakdown of the cell type composition within the tissue. We identified a total of 42,985 cells of which 28,185 (65.6 percent) are PanCK positive tumor cells, 5,917 (13.8 percent) are CD68 positive macrophages, and 8,883 (20.7 percent) are dual negative non-tumor and non-macrophage (neither) cells (**Figure 8A**) across all of the ROIs, with 3,693 cells (8.6 percent) expressing PD-L1 (**Figure 8B**). **Figure 8C** reveals that 41.17 percent of all macrophages are positive for PD-L1. Tumor cells and neither cells are primarily negative for PD-L1, with just 3.31 percent and 3.65 percent positivity for PD-L1, respectively.

**Figure 8.**
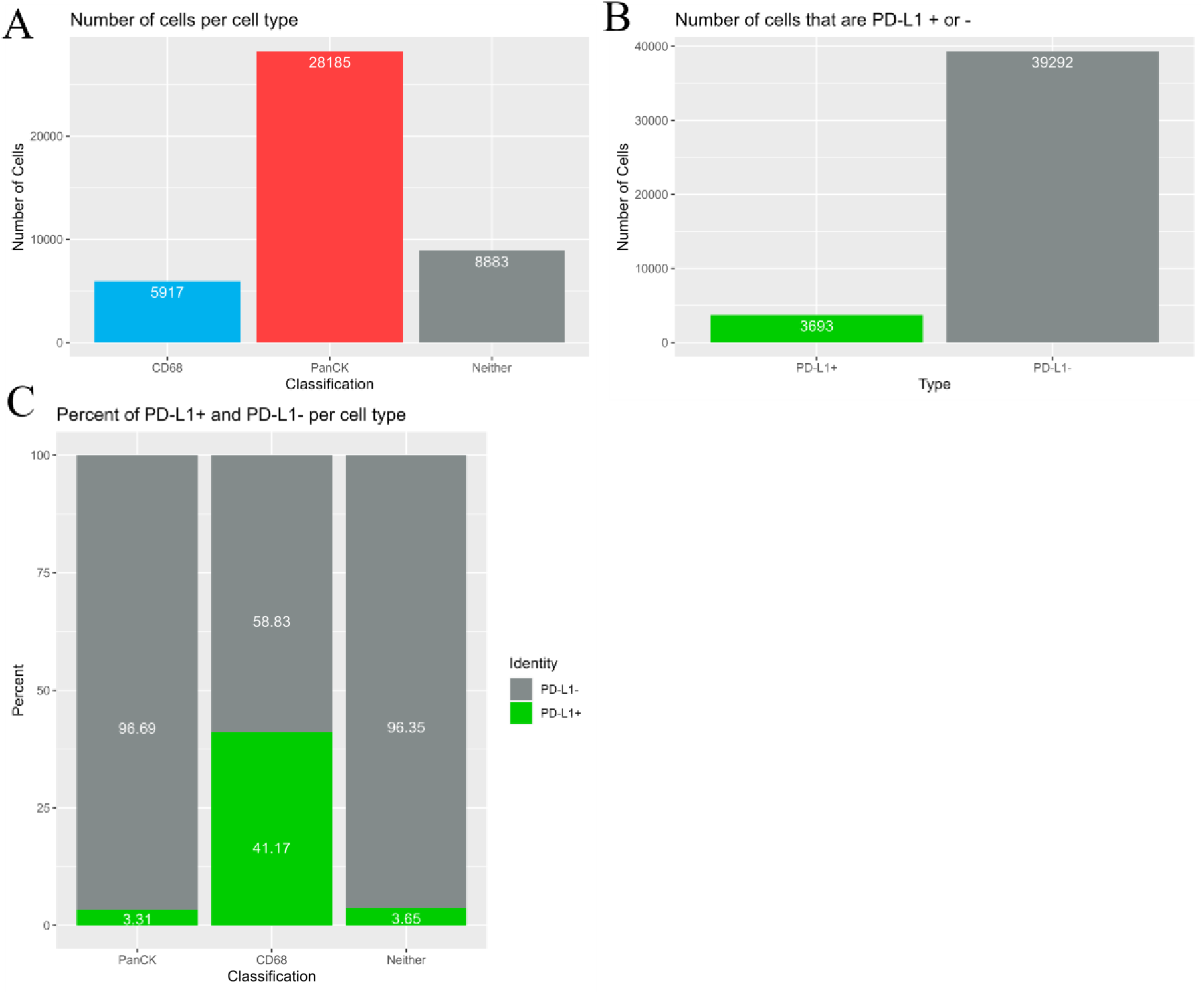
Bar charts displaying summary data for the number of cells for each classification and PD-L1 positivity. **A)** Bar chart showing the number of cells that are macrophages (CD68+), tumor cells (PanCK+), and non-tumor and non-macrophage cells (CD89-/PanCK-). **B)** Bar chart showing the number of cells that are PD-L1 positive and negative. **C)** Bar chart showing the percentage of macrophages (CD68+), tumor cells (PanCK+), and non-tumor and non-macrophage cells (CD89-/PanCK-) that are PD-L1 positive and negative stratified by cell type. The green shading represents the percentage of cells that are PD-L1 positive, and the grey shading represents the percentage of cells that are PD-L1 negative.

This analysis enables the identification of the TME composition through marker recognition and cell segmentation, while also providing us, in this specific example, with primary molecular resolution insights into the immunosuppressive environment.

### 3.2. Single Cell Quantitative Spatial Mapping of Neighborhood Composition and Cell Type Proportions in the PDAC TME

While the initial step of the analysis above provides information on cell typing, it does not address the spatial distribution of these cell types. Understanding the spatial cellular distribution within the TME is essential for deciphering the cellular interactions that contribute to tissue function or pathology. It helps identify potential hotspots of immune activity that can be clinically targeted, enhances the understanding of disease mechanisms, guides therapeutic strategies, and aids in assessing the efficacy of interventions. In this study, a 55 µm radius was defined as the immediate neighborhood surrounding each cell. Within this radius, the spatial coordinates of all neighboring cells were recorded to decipher and quantify direct spatial interactions. This radius, however, can be easily adjusted in the code to suit study specificities. **Figure 9** displays the average proportions of various cell types within a 55 µm radius of each cellular classification. Specifically, **Figure 9A** illustrates the distribution of PanCK^+^ tumor cells within a 55 µm radius from each different cell type. On average, 61.46 percent of the neighboring cells within a 55 µm radius of macrophages are tumor cells, compared to only 46.34 percent for dual negative cells. Furthermore, 74.07 percent of the neighboring cells around tumor cells are also tumor cells. This suggests that tumor cells are more likely to cluster together, potentially creating a more supportive microenvironment for enhanced tumor growth and survival by facilitating cellular communication and evading immune surveillance. When focusing on macrophages (CD68^+^ cells) as neighbors, 17.33 percent of the neighboring cells around macrophages are also macrophages, compared to 11.96 percent for tumor cells and 15 percent for dual negative cells (**Fig. 9B**). Comparing the presence of dual-negative cells (neither) in different neighborhoods, macrophages have 19.46 percent neighboring dual-negative cells, whereas tumor cells have 12.5 percent, and neither cells themselves have 36.08 percent (**Fig. 9C**). Interestingly, even though tumor cells have fewer neighboring macrophages (11.96 percent) compared to neither cells (15 percent) (**Fig. 9B**), their neighborhood includes significantly more immunosuppressive macrophages (PD-L1+/CD68+) than that of neither cells (**Fig. 9D**). Thus, our SNAQ^TM^ platform provides detailed quantitative spatial insights into the immediate neighborhood (defined by a radius of 55 µm from each cell) of a large number of multiple target cell types. To enhance the resolution and refine the stratification of our spatial profiling, we divided the 55 µm radius into three concentric rings (proximal, intermediate, and distal). This approach is based on the concept illustrated in **Figure 1**, with each graph displaying data for a given cell type (**Figure 10**). For instance, **Figure 10A** shows that the proximal neighborhood for tumor cells, on average, consists of 85 percent tumor cells, 9 percent macrophages (with 44 percent of those macrophages being immunosuppressive), and 7 percent non-tumor and non-macrophage cells. In the intermediate ring, the composition changes to 75 percent tumor cells, 13 percent macrophages (with 46 percent being immunosuppressive), and 13 percent non-tumor and non-macrophage cells. The distal ring contains 70 percent tumor cells, 13 percent macrophages (with 46 percent being immunosuppressive), and 17 percent non-tumor and non-macrophage cells, on average. Similarly, our analysis revealed the cellular composition of the three concentric rings surrounding macrophages (**Figure 10B**) and neither cells (**Figure 10C**). The data indicates that, regardless of their relative spatial coordinates, a high proportion of the total non-tumor cells within the TME are macrophages with a significant percentage of them expressing PD-L1, suggesting their immunosuppressive potential. This observation aligns with other research studies reporting a significant proportion of immunosuppressive macrophages within the PDAC TME (Yang, Liu and Liao 2020). Interestingly, the fraction of immunosuppressive macrophages in the tumor cell microenvironment remains relatively constant across the rings, at approximately 45 percent (**Figure 10 A-D**). In contrast, the percentage of immunosuppressive macrophages is significantly lower in the neighborhood of neither cells, ranging from 23.5 percent in the proximal ring, to 29 percent in the intermediate ring, and 33 percent in the distal ring (**Fig. 10C**). This indicates a decreasing gradient of immunosuppressive macrophages as we move closer to the neither cells, whereas the proportion of immunosuppressive macrophages remains consistently higher within a 55 µm radius of tumor cells, regardless of the ring (**Figure 10A**). This suggests that tumor cells attract more immunosuppressive macrophages compared to neither cells. This phenomenon is also observed for macrophages, whose neighborhoods show a high content of immunosuppressive macrophages (47-50 percent) with constant values throughout the three rings (**Figure 10B**). Additionally, when focusing on the microenvironment of immunosuppressive macrophages, we observed a greater proportion of macrophages in their vicinity, with 80 percent, 72 percent, and 66 percent of the macrophages being immunosuppressive across the rings, respectively (**Figure 10D**). These results indicate that immunosuppressive macrophages tend to cluster together or create an environment conducive to their polarization. Thus, our platform allows for quantitative mapping of spatial distribution at a high resolution and provides tailored analysis for detailed and customized investigations.

**Figure 9.**
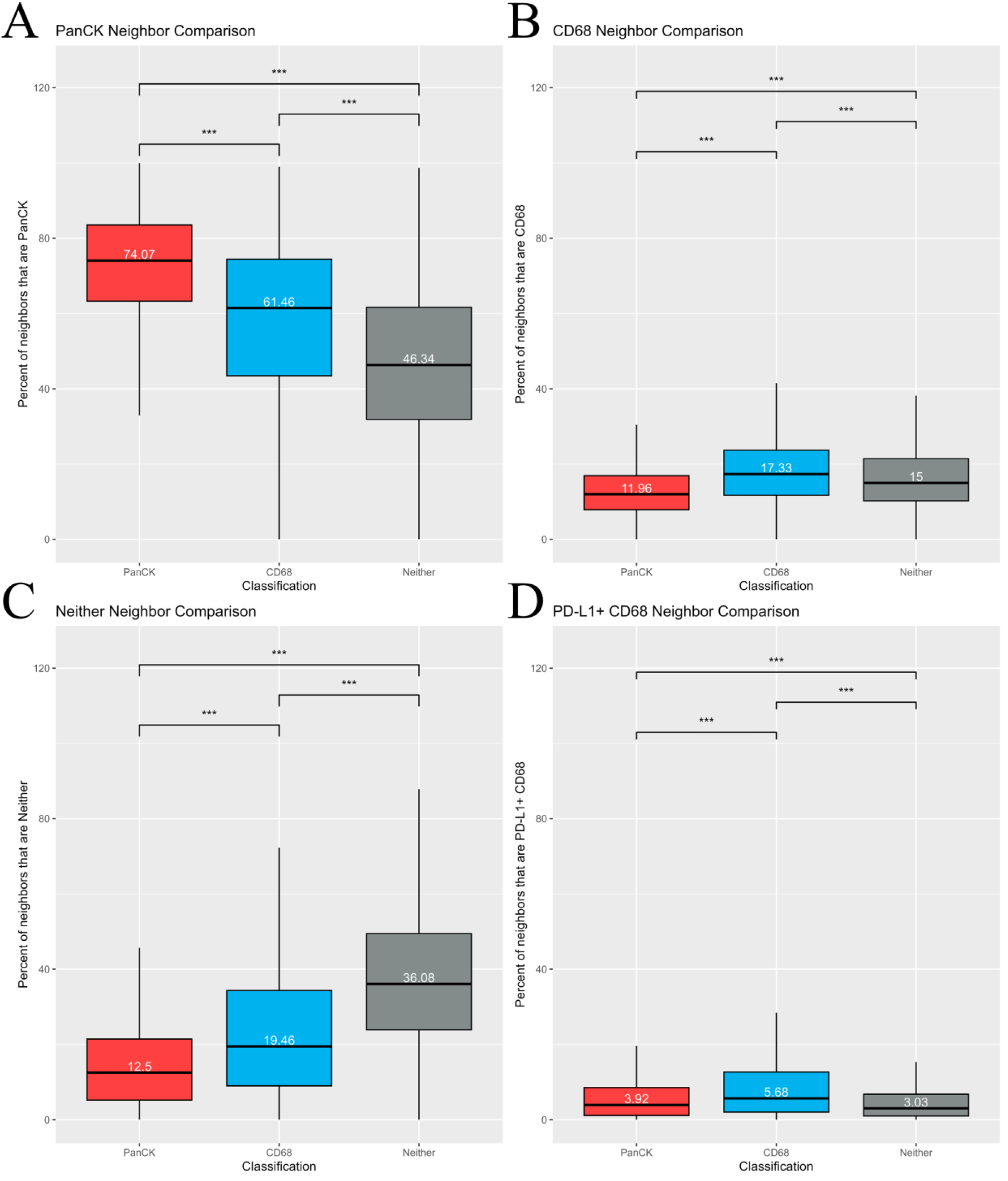
Box and whisker plots showing the percentage of neighboring cell types within the whole 55 µm neighborhood. The average percentage of cells within the 55 µm neighborhood that are tumor cells **(A)**, macrophages **(B)**, neither tumor cells nor macrophages **(C)**, or immunosuppressive macrophages **(D)** are shown for each classification. The bottom and top edges of the boxes represent Q1 (25^th^ percentile) and Q3 (75^th^ percentile), respectively. The height of the boxes represents the interquartile range (IQR). The bold middle line within the boxes represents the median, which is detailed within each box. The bottom whisker extends to the lesser between the minimum data point or the value calculated by (Q1 – 1.5*IQR), and the top whisker extends to the greater between the maximum data point or the value calculated by (Q3 + 1.5*IQR).

**Figure 10.**
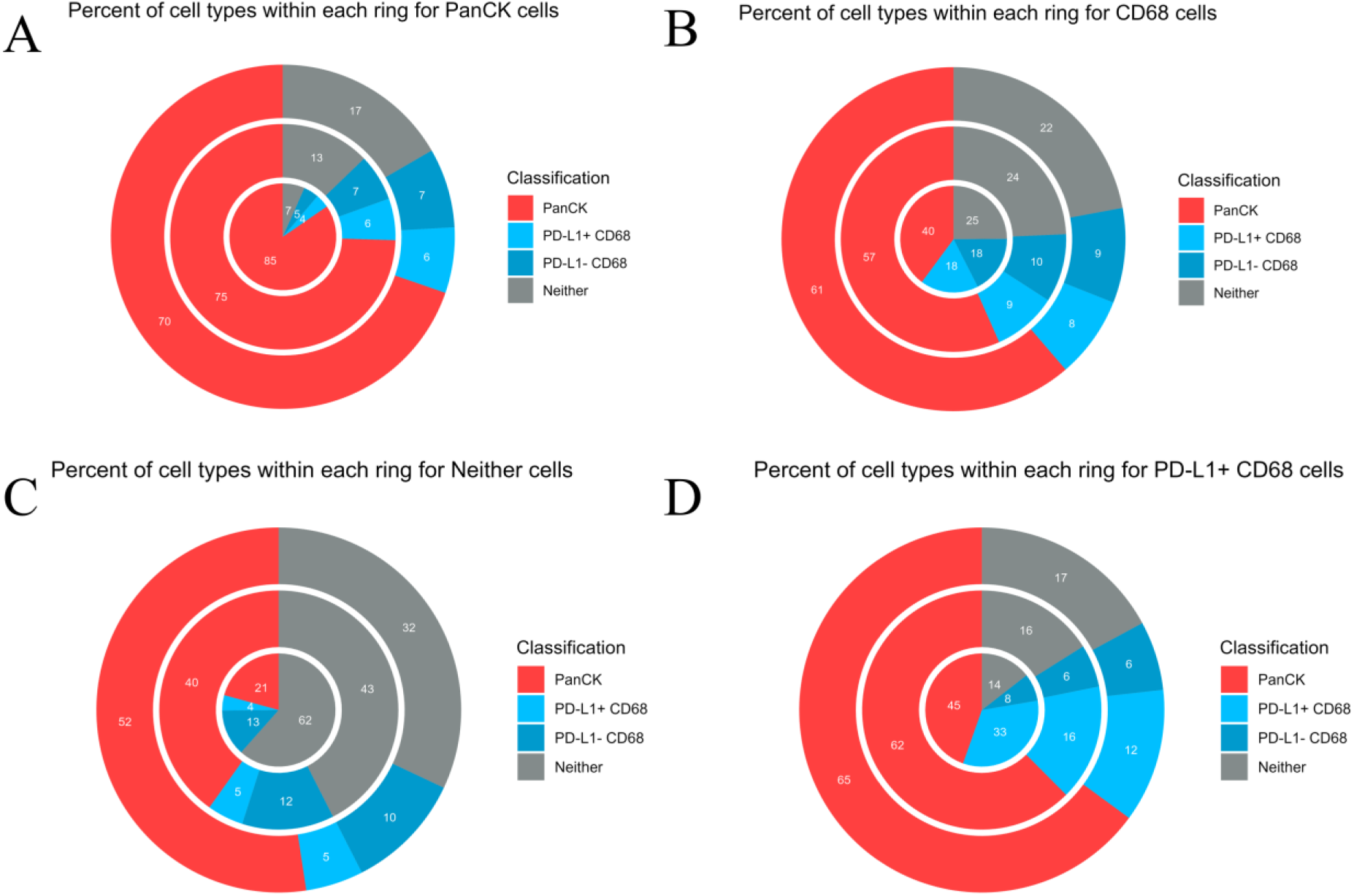
Radial bar charts showing the percentage of the different cell types within each ring. Ring 1 is the central ring, Ring 2 is the middle ring, and Ring 3 is the outermost ring. Panel A measures data derived from only tumor cells, panel B is only macrophages, panel C is only non-tumor and non-macrophage cells, and panel D is only immunosuppressive macrophages.

### 3.3. Closest/Nearest Neighbor Distance Analysis

Analyzing the shortest distance from specific cell types to target cells can provide us with another dimension to better understand the cell types undergoing potential interactions with the target cells. These patterns may define specific functional relationships, potentially indicating particular cell-cell interaction, tissue architecture, active biological processes, disease progression, or response to treatment. This data coupled with the previous data of neighborhood composition and spatial-quantitative mapping of the TME may act as important data for deciphering the interacting partners and their possible biological role. Our study compared the average distance from a macrophage to the closest tumor cell or neither cell. Macrophages find themselves in closer proximity to tumor cells than neither cells (**Figure 11A**). When focusing on immunosuppressive macrophages, this difference is even greater (**Figure 11B**). These observations align with the results presented in **Figure 10** and further highlight the close relationship between tumor cells and immunosuppressive macrophages. The proximity of these macrophages to tumor cells may facilitate immune evasion and promote tumor growth. Interestingly, our study calculated the distance between tumor-associated macrophages and tumor cells in PDAC to be 10.73 µm (**Figure 11A**), and Matusiak and colleagues reported this distance to be 10.6 µm in their analysis of colon and breast tumors using co-detection by indexing (CODEX) computational pipelines (Matusiak *et al*. 2023). This helps demonstrate the validity and reliability of SNAQ^TM^ for single-cell spatial analysis. Furthermore, the similarity also suggests that there might be a conserved spatial relationship across different tumor types, which could be crucial for advancing our understanding of tumor biology and developing targeted therapies.

**Figure 11.**
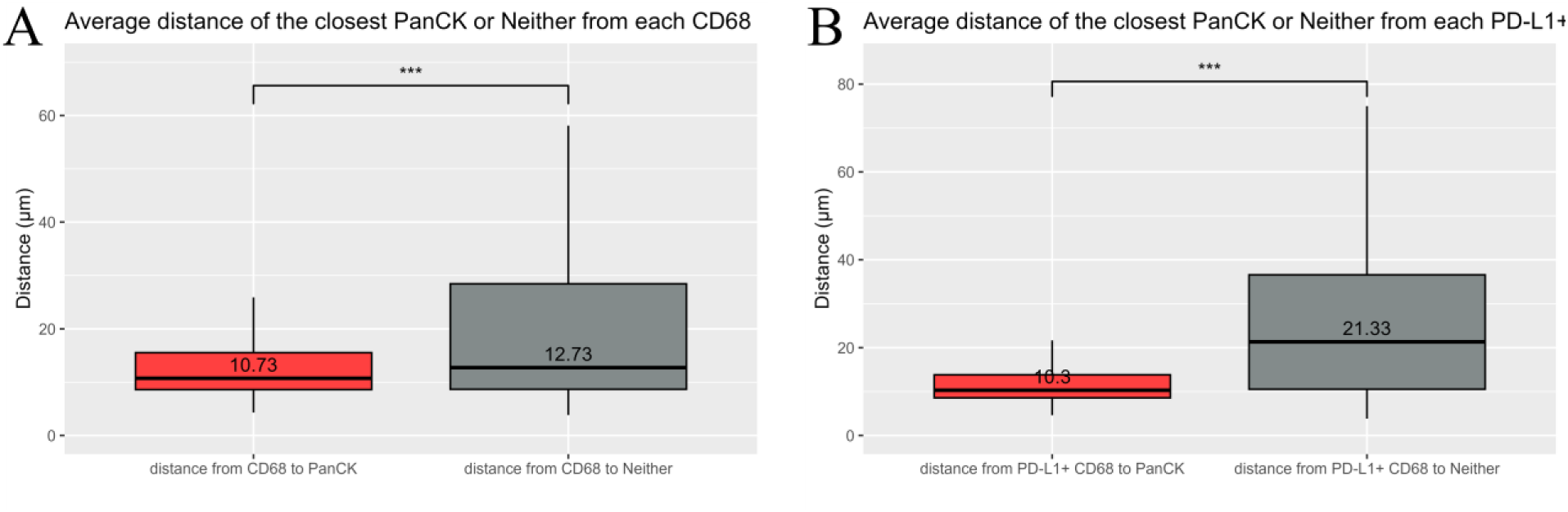
Macrophage-focused cellular closest neighbor distance analysis. **A)** Shows the average distance from each macrophage to the closest tumor cell (red shading) and to the closest non-tumor and non-macrophage cell (grey shading). **B)** shows the average distance from each immunosuppressive macrophage to the closest tumor cell (red), and to the closest non-tumor and non-macrophage cell (grey).

**Figure 12** illustrates the average distance of the closest macrophage or immunosuppressive macrophage from either a tumor cell or a neither cell. Interestingly, tumor cells tend to be farther from the nearest macrophage compared to neither cells (**Fig. 12A**). However, this pattern is reversed when considering immunosuppressive macrophages (**Fig. 12B**). This indicates that immunosuppressive macrophages may have a greater migratory affinity for tumor cells over non-regulatory macrophages, potentially playing a crucial role in facilitating immune evasion and promoting tumor growth. Of note, the differing observations in **Figure 11A** and **Figure 12A** can be explained by the higher concentration of tumor cells relative to macrophages and neither cells in the tissue sample (**Figure 8A**). Given the abundance of tumor cells, each macrophage is likely to have a nearby tumor cell. Conversely, many tumor cells may not have a macrophage within close proximity due to the lower number of macrophages.

**Figure 12.**
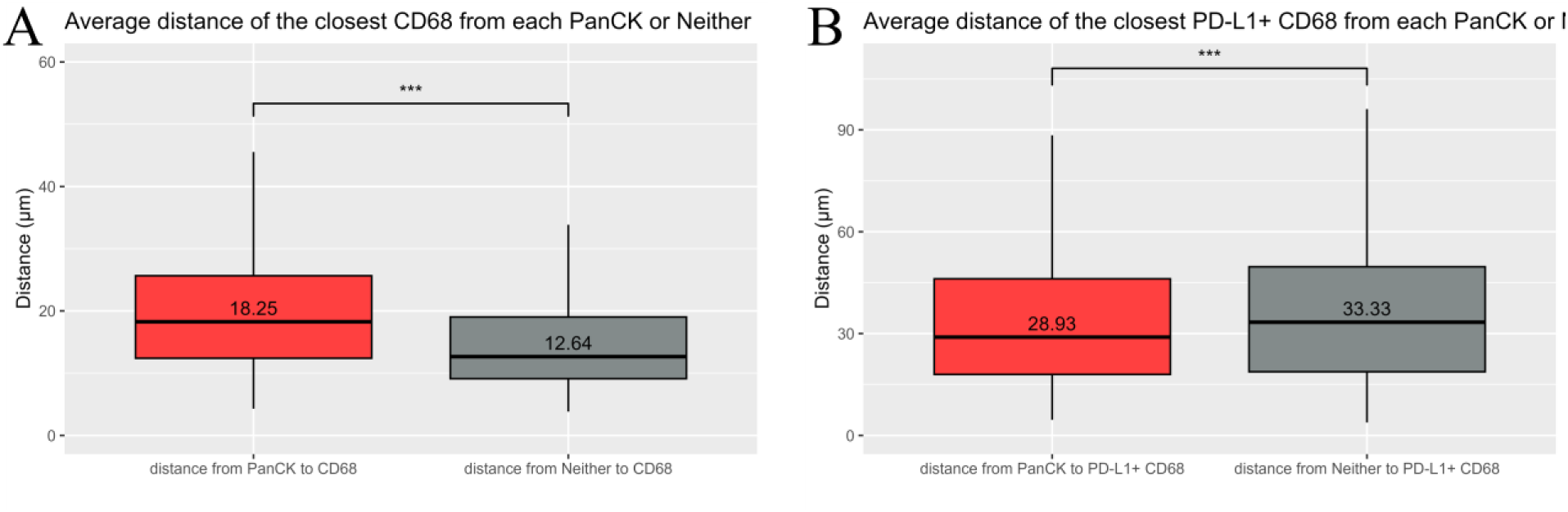
Non-macrophage focused cellular closest neighbor distance analysis. **A)** Average distance from each cancer cell to the closest macrophage (red) from each non-tumor and non-macrophage cell to the closest macrophages (grey). **B)** Average distance from each cancer cell to the closest immunosuppressive macrophage (red), and from each non-tumor and non-macrophage cell to the closest immunosuppressive macrophage (grey).

## 4. Discussion

The SNAQ^TM^ algorithm, which consists of the `Data Analysis.Rmd` and `Plot Maker.Rmd` R Markdown files, is highly customizable to accommodate the variable demands of each user. The platform’s flexibility allows for the adaptation of the algorithm to meet specific analytical requirements. The files can be found in a GitHub repository named “SNAQ” (https://github.com/AryehSilver1/SNAQ). Detailed instructions for modifying the code are found in the ReadMe file. Additionally, the repository includes the input data used in this paper to facilitate code testing and allow for inquiries into the algorithm. This study introduces a free algorithm designed to perform neighborhood analysis on immunofluorescent images. Interrogating tissues with our platform can facilitate better understanding the cell-type distribution and proximities, thereby providing a better understanding of the TME and the potential cellular interactions. Although the algorithm was initially developed for cancer images, it can be applied to any tissue type with the use of markers as indicated in the “Marker Meaning” column in **Table 1**. The SNAQ^TM^ algorithm can offer significant advancements in the field of immuno-oncology by uncovering specific geospatial patterns within tumors, enhancing our understanding of the TME and its interaction with the immune system. By analyzing neighborhood interactions, the algorithm can reveal variations between different patients or treatment groups, highlighting potential differences in patient responses or treatment efficacy. Altogether, this analysis demonstrates that our versatile and comprehensive platform SNAQ^TM^ can effectively dissect and reveal specific heterogeneous cellular patterns and architectures dependent on the cell types composing a tissue. This highlights the power of our method to uncover specific cellular compositions, relationships, and interactions in the context of both normal and malignant tissues. These insights can reveal underlying physiological processes, disease progression, or treatment response. By providing a detailed spatial analysis of cell types and their interactions, SNAQ^TM^ offers a powerful tool for understanding the complexities of the TME. For instance, identifying clusters of immunosuppressive macrophages in proximity to tumor cells can help elucidate mechanisms of immune evasion and inform strategies for enhancing immunotherapy. Similarly, understanding the spatial dynamics of different cell types within normal tissues can shed light on homeostatic processes and identify potential early markers of disease. Furthermore, SNAQ^TM^ promotes data reproducibility by minimizing reliance on manual analysis, thereby helping to standardize results across various studies and ensuring consistency in research findings.

One limitation of the currently presented algorithm is that it can only differentiate between three cell types (letter codes A, B, and D), and the latter is derived through exclusion. However, the code can be modified to accommodate an unlimited number of cell types and functional markers. Since the R Markdown scripts are publicly available, users have the flexibility to edit the code to incorporate any number of markers, tailoring the analysis to their specific research needs.

The versatility of this algorithm extends its utility beyond oncology, making it applicable to a variety of biological and medical research fields. The ability of SNAQ^TM^ to reveal detailed and specific cellular patterns makes it a valuable asset for researchers and clinicians alike. In clinical settings, it can support personalized medicine approaches by providing insights into how individual patients’ tumors might respond to specific treatments based on their unique cellular architecture.

In summary, the SNAQ^TM^ algorithm not only enhances our understanding of cellular heterogeneity and spatial organization within tissues, but also holds significant potential for advancing the diagnosis, prognosis, and treatment of various diseases. The algorithm not only supports cancer research but also holds potential for broader applications in studying various tissue types. Its availability as a free tool encourages widespread use and adaptation, fostering advancements in spatial analysis and contributing to our understanding of complex biological systems.

## 5. Key

Tumor cell = PanCK^+^ cell

Macrophage = CD68^+^ cell

Non-tumor and non-macrophage cell = PanCK^-^ and CD68^-^ cell= Neither = Neither cell

Immunosuppressive macrophage = CD68^+^ and PD-L1^+^ cell

## 6. ACKNOWLEDGMENTS

The PDAC images used for this study were extracted from Aleynick et al. (Aleynick *et al*. 2023) and released under a Creative Common CC BY 4.0 license (https://creativecommons.org/licenses/by/4.0/). This research was partially funded by the National Institute of Health (1R01NS121075, 1R21NS116578, 1R21CA282979 to LPD).

